# Transcriptomic analysis of PER and LEC regions in mice

**DOI:** 10.1101/2025.03.18.643907

**Authors:** Beatriz B. Aoyama, Saúl Lindo-Samanamud, João P. D. Machado, Henrique N. de Araújo, Maria C. P. Athie, André Schwambach Vieira

## Abstract

The perirhinal cortex (PER) is crucial for object recognition memory and comprises areas 35 and 36. The lateral entorhinal cortex (LEC), subdivided into dorsolateral (DLE) and ventral intermediate (VIE) regions, plays a key role in encoding representations of objects, events, contextual details, and sequential information. Despite their anatomical distinctions, gene expression profiling of these subregions remains limited. To address this gap, we conducted transcriptome analysis of these regions in C57BL/6 mice. Tissue samples were collected using laser microdissection, isolating PER areas 35 and 36, as well as DLE and VIE subregions of the LEC, and subsequently analyzed via RNA sequencing. Our results revealed no significant gene expression differences between PER areas 35 and 36, yet substantial molecular divergence emerged when comparing PER to LEC subregions. The PER exhibited higher expression of genes related to nutritional state sensitivity compared to the LEC, while the DLE showed elevated expression of genes implicated in axonal guidance, neurotrophic support, and glutamatergic neurotransmission relative to the VIE. These findings advance our understanding of the molecular profiles unique to PER and LEC regions, and may support the generation of novel hypotheses regarding functional roles of these cortical populations that may be explored in future research.

## 1. INTRODUCTION

In mice, the perirhinal cortex (PER) is anatomically positioned ventrally to the temporal association cortex (TeA) (Beaudin et al. 2013), rostrally to the postrhinal cortex (POR), and dorsally to the lateral entorhinal cortex (LEC) (Agster and Burwell 2009);(Beaudin et al. 2013). Functionally, in rodents, the perirhinal cortex (PER) receives inputs mainly from olfactory, somatosensory, and auditory areas, forming associations that are critical for object recognition, and it also contributes to processing the motivational significance of objects, particularly in approach–avoidance decisions (Buffalo et al. 1999); (Miyashita 2019); (Fiorilli et al. 2021); (Dhawan et al. 2023). Anatomically PER is subdivided into two distinct regions: areas 35 and 36 (Fig. 1). PER area 35 is characterized as agranular, primarily due to the absence of layer VI (Hyman et al. 1986);(Augustinack et al. 2013);(Fiorilli et al. 2021). Area 35 lies ventrally to area 36, dorsally to the piriform area and endopiriform nucleus (rostrally) and entorhinal cortex (caudally) and situated below the rhinal fissure at rostral levels. In contrast, PER area 36 is classified as dysgranular, a key cytoarchitectonic feature distinguishing it from area 35 (Beaudin et al. 2013). Anatomically, PER 35 and 36 are rostrally bordered by Agranular Insular Area, posterior part (AIp) (*Allen Institute for Brain Science, 2004;(Beaudin et al. 2013)*

**Figure 1.**
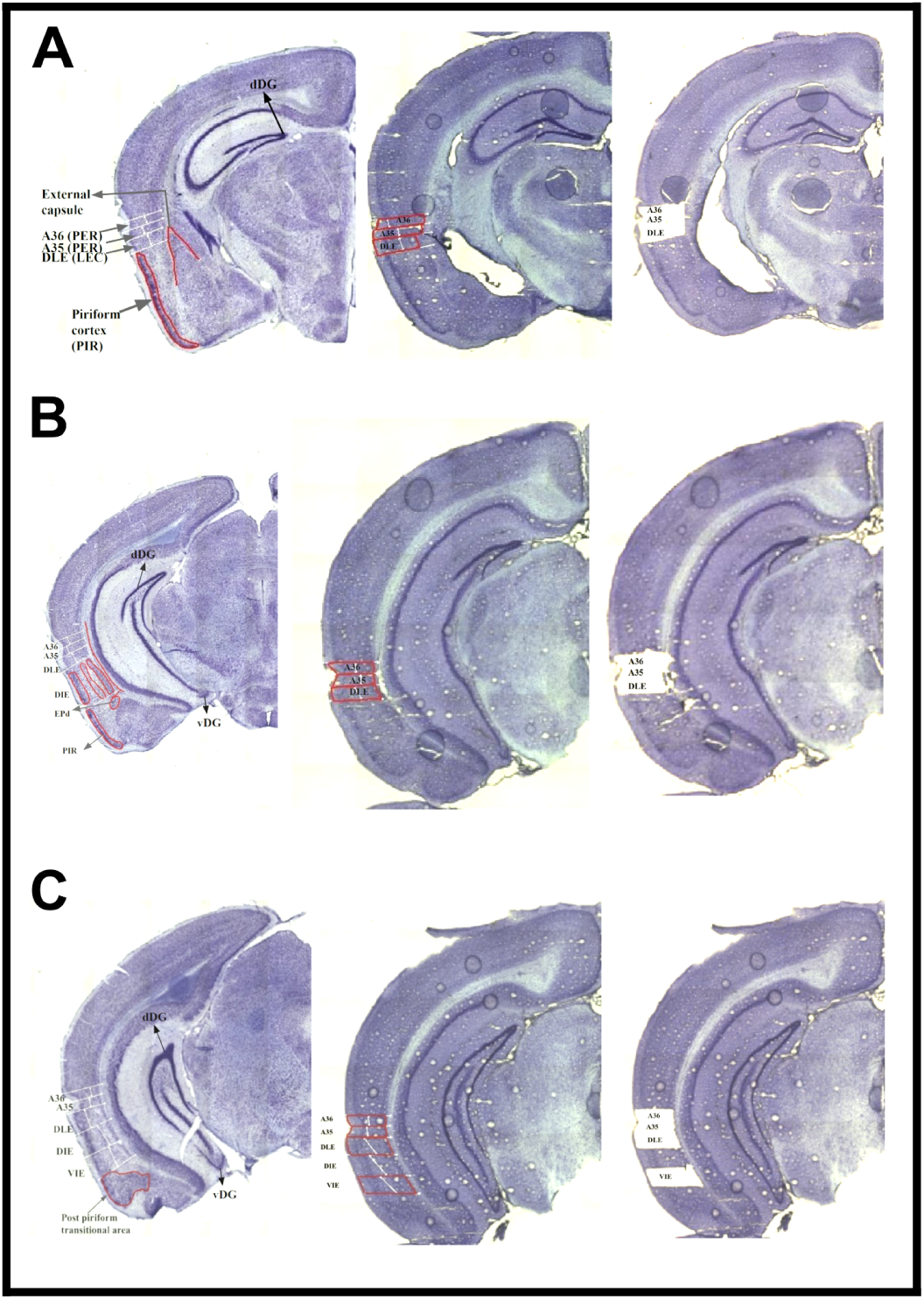
Rostral, intermediate and caudal coronal mouse brain slices for subregions identification showing structures as the dorsal and ventral dentate gyrus, external capsule, piriform cortex and post-piriform transitional area were used as reference to identify the areas 35/36 of PER and subregions of LEC. **(A)** Rostral coronal slices with the PER, DLE and VIE before (Red) and after the LCM. **(B)** Intermediate coronal slices with the PER, DLE and VIE before (red) and after the LCM. **(C)** Caudal coronal slices with the PER, DLE and VIE before (red) and after the LCM.

The lateral entorhinal cortex (LEC), a subdivision of the entorhinal cortex (EC), plays a key role in encoding representations of objects, events, contextual details, and sequential information (Deshmukh and Knierim 2011);(Keene et al. 2016);(Pilkiw et al. 2017). Additionally, this region is also crucial for processes underlying memory formation and learning (Liu et al. 2023);(Tozzi et al. 2024). Anatomically, the LEC is organized into three bands, dorsolateral (DLE), dorsal intermediate (DIE) and ventral intermediate (VIE) <u>(van</u> <u>Groen et al. 2003)</u>(Fig. 1). Anatomical connections of the DLE predominantly target the septal and dorsal regions of the hippocampal formation. Conversely, the ventral-intermediate (VIE) band of the LEC projects mainly to the ventral hippocampal formation via the lateral perforant pathway (van Groen et al. 2003), as well as to the ventral subiculum, although its functional role remains unclear.

The PER and LEC are important components of the hippocampal network, acting as key nodes that integrate information from neocortical and subcortical regions involved in object representation, motivational context, and attentional modulation (Willems et al. 2018). This reciprocal exchange of information between PER and LEC is fundamental for generating object-specific contextual representations and supports the encoding of episodic memories within the hippocampus (Willems et al. 2018). Despite their critical roles in limbic circuitry and their implication in major neurological disorders such as Alzheimer’s disease (Vismer et al. 2015);(Dabrowska et al. 2019) and Temporal Lobe Epilepsy (Petrache et al. 2019);(Di Castro and Volterra 2022), these regions remain underrepresented in transcriptome investigations. Among 197 mouse brain regions most frequently analyzed, approximately 75% of gene expression studies focus on only nine (hippocampus, hypothalamus, striatum, cerebellum, brainstem, pons, medulla, thalamus, and substantia nigra), with PER and LEC notably absent from major datasets (Simpson et al. 2021).

A deeper understanding of PER and LEC molecular architecture is thus crucial for elucidating how these regions contribute to normal cognitive function and how their dysfunction may lead to neurological and psychiatric disorders. To address this gap, the present study combines laser microdissection with RNA sequencing to define the distinct transcriptional signatures of PER areas 35 and 36, as well as the dorsolateral (DLE) and ventral intermediate (VIE) subregions of the LEC in the mouse brain.

## 2. MATERIALS AND METHODS

### 2.1 Animals

Three-month-old male mice (C57BL/6) were used in this study, 5 mice were used for RNA-sequencing and 3 mice were used for histological reference slides preparation. Mice were housed in groups of 4 animals, in a 12-h light/dark cycle on a ventilated rack with free access to food and water throughout the experimental process. The experiments were performed at the University of Campinas and the experimental protocol was approved by the University research ethics committee (CEUA 5682-1; 2020 protocol) according to accepted ethical practices and legislation regarding animal research in Brazil (Brazilian federal law 11.794 from October 8th, 2008).

### 2.2 Morphological identification of cortical areas of interest

Areas 35 and 36 of the PER, as well as the DLE and VIE subregions of the LEC, were identified based on The Allen Mouse Brain Atlas (portal.brain-map.org) and previously published studies <u>(van Groen et al. 2003)</u>;(Beaudin et al. 2013). Histological reference slides were produced in a Cryostat using fixed brains. Coronal serial sections of 45 µm were produced for each brain, mounted on glass slides and Nissl stained. Slices were categorized as rostral, intermediate, or caudal based on hippocampal morphology as a landmark. The width parallel to the cortical surface for each region was used to determine the anatomical boundaries between adjacent regions.

Rostral slices were selected between Bregma -1.82 and -2.80. Rostral slices included the granular layer of the dorsal dentate gyrus, external capsule, endopiriform nucleus and piriform cortex as reference structures (Fig.1A). Area 36 was identified based on its lateral distance from area 35, as both form horizontal bands along the rostrocaudal axis, according to the 3D model from the Mouse Brain Atlas. In these sections, the lateral distance measured was 196-197 µm for areas 35 and 36 of the PER, and 288-289 µm for the DLE region. Intermediate slices were selected between Bregma -2.92 and -3.64. The intermediate slices included the dorsal and ventral dentate gyrus, dorsal and ventral CA3, and post-piriform transitional area as reference structures (Fig.1B). In these sections, the lateral distance measure was 196-197 µm for areas 35 and 36 of the PER and 400-401 µm for the DLE region.

Caudal sections were selected between Bregma -3.80 and -4.04. These caudal sections included the same reference structures as the intermediate sections (Fig. 1C), however, for slices near Bregma -4.04, we considered sections without the presence of CA3, but with both dorsal and ventral dentate gyrus, absence of the polymorphic layer, and presence of the medial band of the dorsal zone of the entorhinal cortex. In these sections, the lateral distance measure was 196-197 µm for areas 35 and 36 of the PER, 400-401 µm for the DLE subregion and 411-412 µm for the VIE subregion.

### 2.3 Laser Microdissection

The morphological identification parameters stated above were used to identify areas 35 and 36 of PER, and subregions DLE and VIE of LEC in frozen tissue sections. For laser microdissection, previously frozen brains were sectioned using a cryostat (Leica Biosystems) to obtain 60 µm serial sections for RNA sequencing, with the entire hippocampus used as a reference structure. The coronal slices were collected in PEN membrane-covered slides (Life Technologies®, Thermo Fisher Scientific) and these slides were stained with Cresyl Violet, dehydrated with an ethanol series and stored at -80 °C. The structures were delimited using a PALM (Zeiss®) system and the tissue was mechanically collected in separate tubes using surgical microscope and ophthalmic forceps.

### 2.4 RNA isolation, cDNA library preparation and RNA sequencing

RNA was isolated from microdissected samples using TRIzol’s manufacturer protocol for RNA isolation (Thermo Fisher Scientific). Subsequently, cDNA libraries were produced using NEBNext Poly(A) mRNA Magnetic Isolation module for mRNA isolation and VAHTS Universal V8 RNA-seq Library Prep Kit for MGI kit following the manufacturer instructions. Sequencing was performed in a Novaseq platform (Illumina) available from Macrogen Inc, producing an average of 22 million reads per sample. Reads were aligned to the *Mus musculus* reference genome GRCm38 Ensembl release 102 assembly using the STAR aligner tool. The average unique alignment percentage was 83,14% and for the gene expression estimation and statistical analysis purposes, we use the the DESEq2 statistic package (http://www.bioconductor.org/packages/release/bioc/html/DESeq2.html).

A list of differentially expressed genes with a statistical significance of p < 0.05 (after correction for multiple tests) was generated. This list was submitted to enrichment analysis using clusterProfiler package (https://bioconductor.org/packages/release/bioc/html/clusterProfiler.html, RRID:SCR_016884) (Yu et al. 2012) and the functional profiles were classified into KEGG Pathways.

With the aim of identifying differentially expressed genes and enriched signaling pathways, comparative transcriptomic analyses were performed between areas 35 and 36 of the PER and the DLE and VIE subregions of the lateral entorhinal cortex, analyzed separately. The comparison between areas A36 and A35 revealed only one differentially expressed gene (*Ly86*), which was upregulated in A36. Due to the minimal transcriptomic differences observed between these two areas (Fig. S1), this comparison was excluded from further analyses. Consequently, areas 35 and 36 were grouped and considered together as the PER region in subsequent analyses.

### 2.5 Transcriptomic data cross-validation

The cross-validation involved the selection of *in situ* hybridization (ISH) sections images available on the online platform of the Allen Mouse Brain Atlas (https://mouse.brain-map.org/). We employed the same identifying criteria used for laser microdissection of PER, DLE, and VIE along the rostro-caudal axis in order to select ISH section images. The genes *MC4R, Lepr, Sema3E, Slc17a8, Bdnf*, and *Gfra1* were analyzed, with an average of 17 sections per gene. Using the ImageJ software (Version 1.46r), the regions of interest were selected with the ROI Manager tool, and an average threshold (120-188.16) was established. The average percentage of the area was measured after applying the average threshold to the areas of interest, PER, LEC (DLE and VIE), in which we used the same methods to identify the regions described in the methodology. The average percentage of the area was measured considering rostral, intermediate, and caudal sections, as well as the separation of gene expression according to the rostro-caudal axis.

## 3. RESULTS

### 3.1 Differentially expressed genes identification

Principal component analysis (PCA) demonstrated clustering of the samples according to their respective regions (Fig. 2A). Furthermore, we identified 657 differentially expressed genes in PER vs. DLE, including 248 more expressed in PER when compared to DLE and 409 less expressed in PER when compared to DLE (Fig. 2B). In the PER vs. VIE comparison, we found 1,878 differentially expressed genes, with 848 more expressed in PER when compared to VIE and 1,029 less expressed in PER when compared to VIE (Fig. 2B). Lastly, in the DLE vs. VIE comparison, 968 differentially expressed genes were identified, in which 480 were more expressed in DLE when compared to VIE and 488 were less expressed in DLE when compared to VIE (Fig. 2B). The complete list of differentially expressed genes is available in Table S1. The Venn diagram illustrates that 118 differentially expressed genes are shared among PER vs. DLE, PER vs. VIE, and DLE vs. VIE comparisons (Fig. 2C).

**Figure 2.**
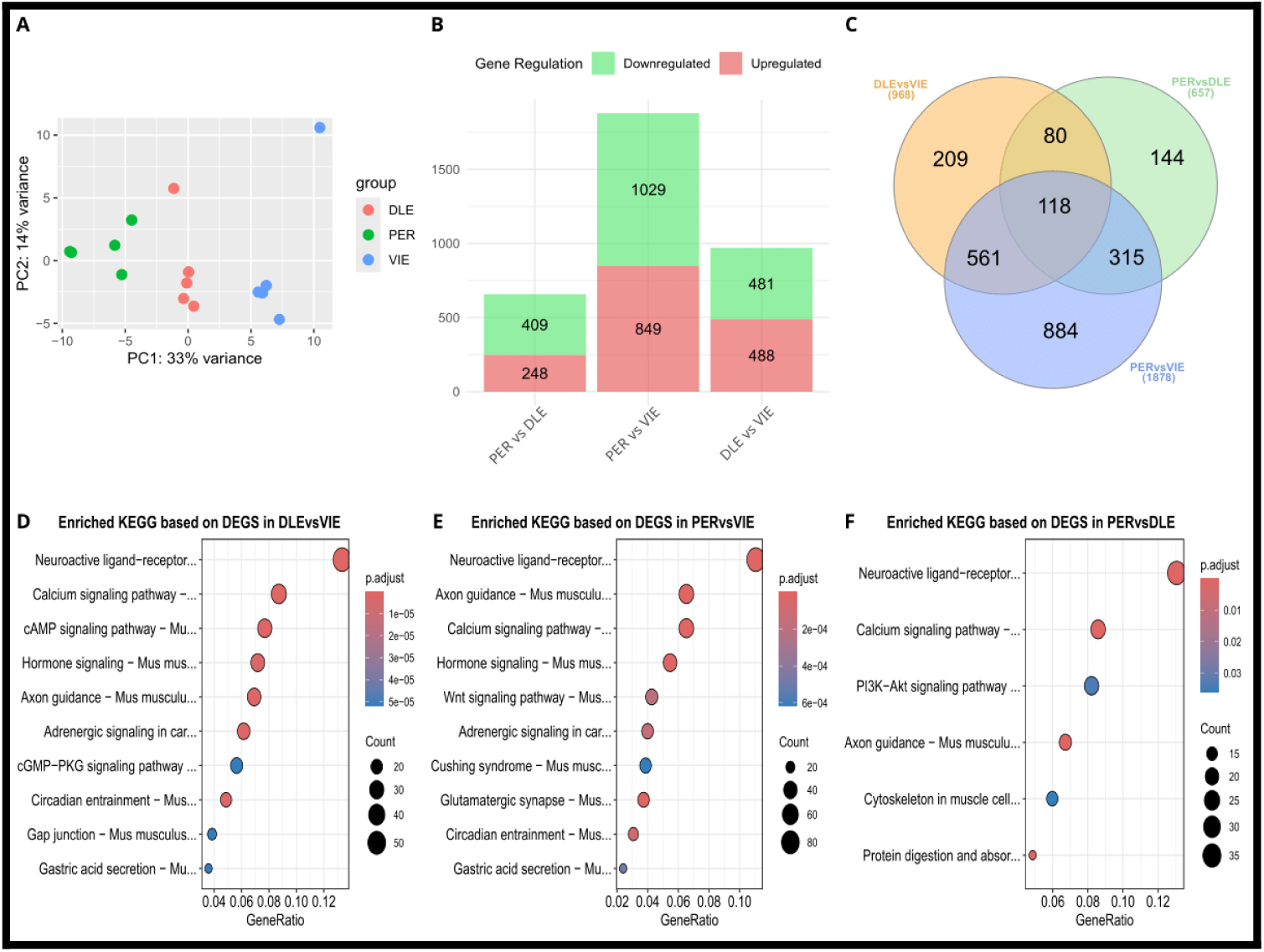
Sample variability and functional enrichment analysis. **(A)** The PCA considered the transcriptomic analysis 2. **(B)** Total number of differentially expressed genes. **(C)** Venn diagram representing common and unique differentially expressed genes. **(D)** A dot plot of DLEvsVIE enriched KEGG pathways. **(E)** A dot plot of PERvsVIE enriched KEGG pathways. **(F)** A dot plot of PERvsDLE enriched KEGG pathways.

### 3.2 Enrichment Analysis

The enrichment analysis using the KEGG database revealed a total of 6 significant enriched pathways (p.adjusted <0.05) for PER vs. DLE (Fig. 2D); 49 significant enriched pathways (p.adjusted <0.05) for PER vs. VIE (Fig. 2E) and 55 significant enriched pathways (p.adjusted <0.05) for DLE vs. VIE (Fig. 2F). For a complete list of KEGG pathways refer to Table S2.

### 3.3 *in silico* ISH cross-validation

To cross-validate our findings, we selected key genes from the most enriched pathways and explored ISH data from the online Allen Mouse Brain platform. The genes *Lepr, MC4R, Sema3E, Bdnf, Slc17a8*, and *Gfra1* (Fig. 3A), were selected for such validation. We measured the percentage of area along the hippocampal rostro-caudal axis in the regions of interest (PER, LEC, DLE, and VIE) according to gene expression patterns. *MC4R* ISH data showed an average area percentage of 3.1 in PER and 1.2 in LEC (Fig. 3B), supporting our hypothesis of higher gene expression in the PER compared to the DLE and VIE of the LEC. Additionally, *MC4R* expression was highest in intermediate and caudal sections of the PER compared to the same sections in the LEC. For the *Lepr* gene, our data showed an average area percentage of 1.4 for PER, 1.9 for DLE, and 1 for VIE. However, unlike *MC4R*, the highest expression of the *Lepr* gene by ISH was present in rostral sections in PER and DLE.

**Figure 3.**
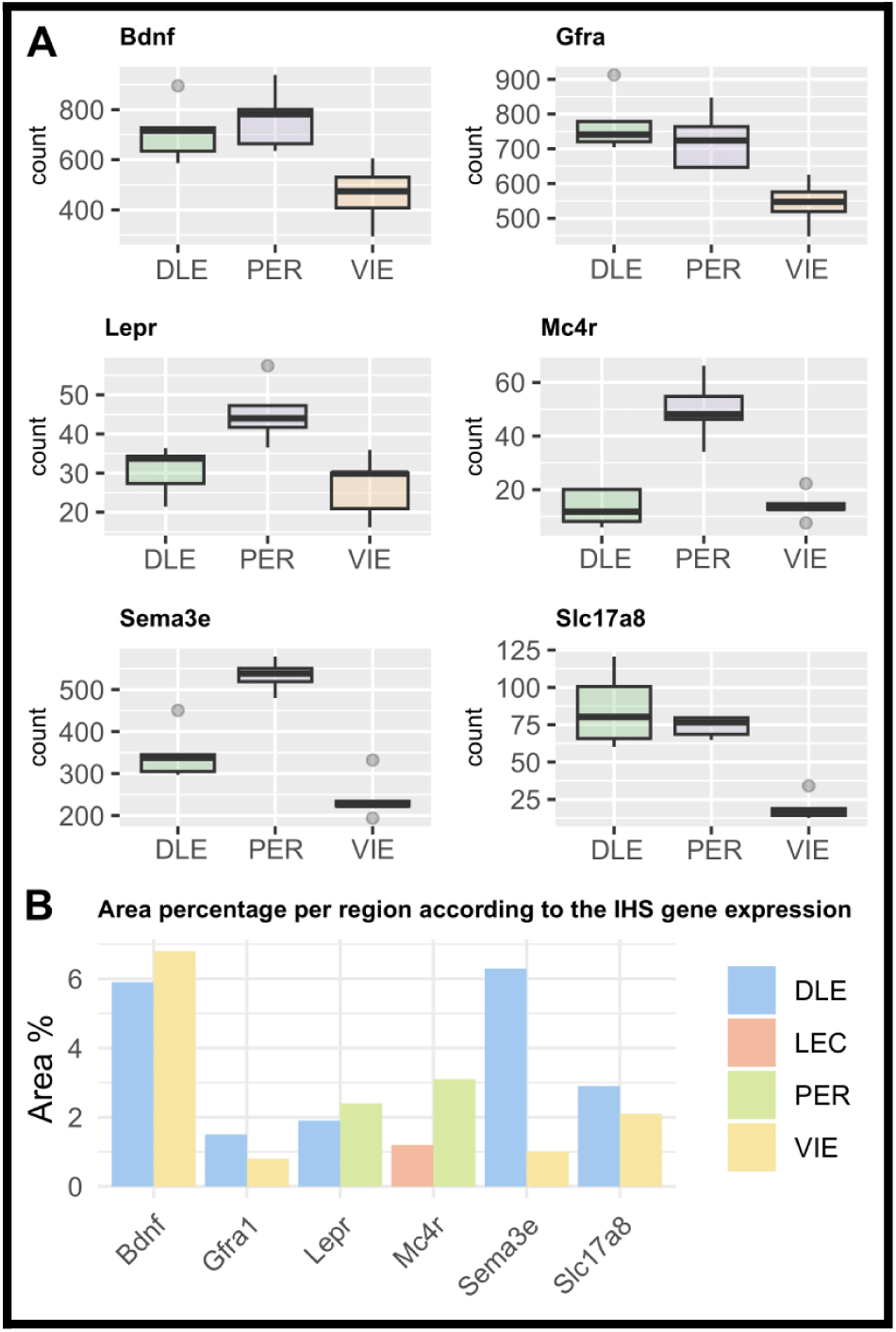
In situ hybridization (ISH) for differentially expressed genes in PER, DLE and VIE. **(A)** Estimated gene expression level graphic for each subfield. **(B)** Area percentage per region according to the gene expression of IHS for the possible marker genes.

In the validation of genes associated with axon guidance, synaptic neuroplasticity mechanisms, and glutamate transport in the comparison of DLE with VIE, we observed an average area percentage of 6.3 in DLE and 4.5 in VIE for the *Sema3E* gene (Fig. 3B). ISH data indicated that *Sema3E* was more highly expressed in caudal sections of DLE. We also observed higher expression of the *Slc17a8* and *Gfra1* genes by ISH in DLE (1.9 and 1.5, respectively) compared to VIE (2.1 and 0.8, respectively) (Fig. 3B). However, *Slc17a8* expression was highest in intermediate and caudal sections of DLE, while *Gfra1* expression was concentrated in rostral sections of DLE. Regarding the *BDNF* gene, we observed an average area percentage of 6.8 in the VIE and 5.9 in the DLE (Fig. 3B).

## 4. DISCUSSION

This study presents for the first time the transcriptomic profile of mice’s PER and the subregions of DLE and VIE of LEC obtained through laser capture microdissection (LCM). Our results indicate that the PER subregions display highly similar transcriptional profiles, most likely attributable to their spatial proximity. On the other hand, physical distance of VIE from PER and DLE may have an influence in the higher number of differentially expressed genes and pathways found in the pairwise comparison to these regions (Yao et al. 2023). Notably, all comparisons (PER vs. DLE, PER vs. VIE, and DLE vs. VIE) converged on a common enriched pathway in the KEGG database, the ‘neuroactive ligand-receptor interaction.’ This pathway integrates multiple signaling mechanisms mediated by distinct classes of neural receptors. However, the specific receptor-related pathways contributing to this enrichment differed across each comparison, suggesting region-specific signaling dynamics within a shared broader functional framework.

The observed higher expression of genes that code for the receptors of melanocortin, glucagon and leptin in PER compared to LEC suggests that PER may be more responsive to hormonal signals related to food-associated object recognition. These increased sensibility to hormone signaling would activate limbic circuits involved in memory and learning processes that support food-seeking and feeding behaviors. When comparing DLE to VIE, the higher expression of genes associated with axon orientation, neurotrophins and production and vesicular transport of glutamate indicates an enhanced neuroplasticity machinery in DLE when compared to PER or VIE.

### 4.1 Nutritional state sensitivity in PER vs. LEC

Building upon our transcriptomic findings, we examined genes encoding receptors involved in metabolic signaling, including leptin, melanocortins, and glucagon-like peptides, to explore the functional implications for PER and LEC. We observed a higher expression of the *MC4R* gene in PER (areas 35 and 36) relative to LEC, the *GLP2R* gene in PER (areas 35 and 36) relative to LEC, and the *Lepr* gene in PER (area 36) relative to VIE. These results were cross-validated using in situ hybridization (IHS) data from the Allen Mouse Brain Atlas, which confirmed higher expression of these genes in PER compared to LEC and VIE.

Leptin, secreted by adipocytes, signals metabolic state to the brain, with its receptor expressed in hypothalamus, cortex, amygdala, hippocampus, brainstem, and cerebellum (Harvey 2007). MC4R participates in the hypothalamus–leptin–melanocortin system controlling feeding and energy homeostasis (Tao 2010) (Li et al. 2019). GLP2R, the receptor for glucagon-like peptide-2, is involved in feeding control and neuromodulation (Lovshin et al. 2001); (Amato and Mulè 2019). Together, these findings suggest that PER neurons may be more responsive than LEC neurons to multiple metabolic signals.

Importantly, PER and LEC neurons expressing long-form leptin receptor (ObRb) project to the hypothalamic arcuate nucleus (DeFalco et al. 2001), providing anatomical support for metabolic input. The higher expression of *Lepr*, *MC4R*, and *GLP2R* in PER may reflect a functional integration linking internal nutritional state with higher-order cognitive processes. In ecological terms, this could support survival behaviors by connecting memory circuits to feeding: recognition of objects (potentially food), their locations, palatability, and familiarity would activate PER-dependent memory and learning pathways during exploration and foraging (Higgs and Spetter 2018);(Seitz et al. 2021).

Overall, our data suggest that PER, relative to LEC, is particularly sensitive to hormonal signaling associated with energetic state, implying that fluctuations in nutritional status could modulate PER function and related cognitive processes. Future functional studies are required to directly test these hypotheses.

### 4.2 Axon guidance and neuroplasticity genes in DLE vs. VIE

Our data also evaluated genes associated with axonal guidance, neurotrophic factors, glutamate production and transport, which suggest possible pro-excitatory synaptic plasticity mechanisms in the comparison between DLE and VIE. We observed higher expression of *Sema7A*, *Sema3E*, *Efna5*, *EphA7*, *BDNF*, *GFRA1*, *GLS2* and *Slc17a8* genes in DLE when compared to VIE. Similar to the previous findings, these results were likewise cross-validated using IHS data from Allen Mouse Brain Atlas, which confirmed higher expression of these genes in DLE compared to VIE.

The axon guidance is a dynamic process, in which guidance receptors are expressed in growing ends of motile axons and it interacts with signaling molecules or membrane-anchored molecules, resulting in the attraction, repulsion, or growth of axonal extensions (Accogli et al. 2020). This guidance process is important for remodeling, maintenance and organizing mature synapses(Yuasa-Kawada et al. 2023). Sema3E is part of the semaphorin class 3 family and it binds to neuropilin and plexin receptors, being an important component of the axon guidance process (Takahashi et al. 1999). This interaction is critical for mediating repulsive signals in GABAergic interneurons in deep layers Vb and VI of primary motor area, secondary and primary somatosensory area and primary visual area in mice(Polleux et al. 2000);(Watakabe et al. 2006). Sema7A interacts with integrin receptor β1, this interaction is important to promote dendritic growth in mature granule neurons in dentate gyrus in mice adult neurons(Jongbloets et al. 2017).

Efna5 is expressed in adult mice ‘cortex and it binds to the Epha7 receptor. This signaling molecule is expressed in the entorhinal cortex, mainly in layers 2,3 and 5 (Cooper et al. 2009), as well as granular neurons of the dentate gyrus (Hara et al. 2010). In granular neurons of the dentate gyrus, the knockdown of the Efna5 gene compromises the maintenance and proliferation of newly formed granular neurons. The deletion of the *EphA7* gene (EphA7−/−) in developing cortical neurons followed by ephrin-A5 exposuring surface in vitro, resulted in decreased repulsion, increased growth of dendritic spines and filopodia during postnatal development, and decreased excitatory synapse formation compared to wild-type neurons (Clifford et al. 2014).

BDNF is a neurotrophic factor widely expressed in both excitatory and inhibitory synapses, having a dependent activity related to synaptic plasticity (Tyler et al. 2002);(Wardle and Poo 2003). Studies have been showing the mature form of BDNF and kinase tyrosine (TrkB) receptor are widely expressed in hippocampus and entorhinal cortex in rats (Yan et al. 1997);(Kaplan and Miller 2000), this interaction is important in synaptic plasticity mechanism in learning (Yan et al. 1997);(Kaplan and Miller 2000), memory recognition, and spatial memory (Kesslak et al. 1998);(Mizuno et al. 2000);(Cirulli et al. 2004);(Bekinschtein et al. 2007);(Heldt et al. 2007). The activation of TrkB by BDNF stimulates exocytosis of synaptic vesicles containing glutamate at presynaptic sites by activation of voltage-gated calcium channels (VGCC) (Leal et al. 2017).

GFRα1 is a receptor of GDNF (glial cell line-derived neurotrophic factor) and it is expressed in the cortex, olfactory areas and hippocampus formation (Trupp et al. 1997). This interaction is important in dendritic spines and dendrites development in excitatory hippocampal neurons in vivo and in vitro (Irala et al. 2016).

Although the entorhinal cortex is known to contribute to synaptic plasticity due to its role as a transitional region between the hippocampus and other cortical areas (Buzsáki, 1989), there is a lack of studies investigating neuroplasticity mechanisms along the rostrocaudal axis of the lateral entorhinal cortex. Therefore, our study is the first to present molecular data demonstrating distinct synaptic plasticity mechanisms between the DLE and VIE of LEC.

Taken together, our findings suggest that DLE neurons may have a more abundant molecular machinery associated with pro-excitatory neuroplasticity mechanisms when compared to VIE. This hypothesis could also be supported by the observed higher expression of genes associated with glutamate production and transportation to the synaptic cleft (higher expression of *GLS2* and *Slc17a8* genes) in DLE related to VIE. These observed differences in gene expression may be associated with the different projections from DLE and VIE to the hippocampus formation, but further studies should further explore the functional implications such differences in the molecular machinery of these regions.

## 5. Conclusion

Overall, our data suggest that PER, compared to LEC, is particularly sensitive to hormonal signaling associated with energetic state, implying that fluctuations in nutritional status could modulate PER function and related cognitive processes. On the other hand, the comparison between DLE and VIE demonstrated a more abundant molecular machinery associated with pro-excitatory neuroplasticity mechanisms in DLE compared to VIE. These findings advance our understanding of the molecular profiles unique to PER and LEC regions, providing a detailed characterization of their gene expression landscapes. Furthermore, these data may support the generation of novel hypotheses regarding functional roles of these cortical populations that may be explored in future research.

## 6. Acknowledgments and author contributions

Special thanks to Professor Menno P. Witter for his critical insights and suggestions for the final version of this manuscript, especially regarding the neuroanatomical references mentioned.

## Author contributions

Beatriz B. Aoyama (Formal analysis, Data curation, Visualization, Writing original draft, Writing—review & editing), Saúl Lindo Samanamud (Formal analysis, Visualization, Writing original draft, Writing—review & editing), João P. D. Machado (Data curation, Visualization,Writing—review & editing), Henrique Nascimento de Araujo (Visualization, Writing — review & editing), Maria Carolina Athie (Visualization, Writing — review & editing), André Schwambach Vieira (Conceptualization, Funding acquisition, Visualization, Writing—review & editing)

## 7. Supplementary material

Additional supporting information can be accessed online in the *Cerebral Cortex* supplementary materials.

## Supplementary Information

**Figure S1 Intragroup and intergroup variability**.The figure shows PCA analysis and gene regulation in A35, A36, DLE, and VIE. PCA (A) reveals separation along PC1 (28%) and PC2 (14%), indicating transcriptional differences. Panel (B) displays upregulated and downregulated genes, highlighting distinct expression profiles and unique transcriptional signatures across experimental conditions.

**Supplementary Table 1. Differentially expressed genes.**

**Supplementary Table 2. Enrichment pathways (KEGG).**

**Supplementary Table 3. Most expressed genes in PER, DLE and VIE.**

